# Automated Workflow For Peptide-level Quantitation From DIA/ SWATH-MS Data

**DOI:** 10.1101/2020.01.21.914788

**Authors:** Shubham Gupta, Hannes Röst

**Author notes:** Corresponding Author: Hannes Röst.

## Abstract

Data Independent Acquisition (DIA) is a powerful method to acquire spectra from all ionized precursors of a sample. Considering the complexity of the highly multiplexed spectral data, sophisticated workflows have been developed to obtain peptides quantification. Here we describe an open-source and easy-to-use workflow to obtain a quantitative matrix from multiple DIA runs. This workflow requires as prior information an “assay library”, which contains the MS coordinates of peptides. It consists of OpenSWATH, pyProphet and DIAlignR software. For the ease of installation and to isolate operating system related dependency, docker-based containerization is utilized in this workflow.

## Introduction

Liquid Chromatography coupled to tandem Mass-Spectrometer (LC-MS/MS) is widely used to analyze the proteome of biological samples. Currently, multiple methods using LC-MS/MS are available to practitioners which include targeted proteomics, shotgun proteomics and data-independent acquisition (DIA) [1]. Over the last decade, DIA has gained traction for high-throughput and reproducible analysis [1-3]. Compared to traditional shotgun proteomics, in DIA/SWATH-MS peptides are isolated using a larger m/z window and are co-fragmented, which results in rich multiplexed MS2 spectra from multiple precursors. Experimental guidelines to generate DIA data are explained in Chapter 11 (by Hauck et al). Since DIA spectra are convoluted with fragment-ions from many precursors, innovative strategies have been developed to extract signals for each peptide of interest. These approaches are divided into two categories: library based (peptide-centric) and non-library based (spectrum-centric). In general, library-based methods tend to be more sensitive and produce accurate quantitative results, especially if the assay library is prepared from the same sample [4]. In this chapter, we will focus on the library-based workflow.

Efficient ways to obtain a high-quality library and guidelines are detailed in the article by Schubert et al [5], and in chapter 31 (by Eggers et at). Recent studies suggest that libraries can also be efficiently predicted using computational means alone [6, 7]. A library consists of a collection of MS coordinates uniquely describing the peptides of interest. This includes the elution time of peptides, their charge state, fragment-ions that are most representative (unique and detectable) and their relative intensities. Each precursor is called an “assay” and this assay library is the key for a successful DIA data analysis. In addition, the assay library must include decoy-peptides for statistical scoring [8, 9]. Note the DIA data contains spectra from all ionized precursors which makes it suitable for re-mining with a new library, however proper diligence is needed [10].

Compared to automated workflow for traditional mass-spectrometer data in which spectra are matched against a database, a DIA workflow focuses on identifying the correct chromatographic peak and its coelution profile; similar to targeted proteomics analysis [11]. Firstly, raw spectra files are converted from the vendor-specific format to a standard format such as mzML. For format-conversion, we are using the MSConvert tool [12].

Feature extraction is performed by first aligning library retention time (RT) to spiked-in standards, for example the iRT peptides, followed by fetching MS2 extracted-ion chromatograms (XICs) using library coordinates, as shown in Figure 1. OpenSWATH, supported by OpenMS, is an open-source software which performs this task [11]. OpenSWATH picks multiple peaks from XICs of a precursor, therefore, a statistical method is needed to assign a probability to each peak for being the correct one; in addition, chromatograms without any signal also need to be correctly labelled as such. pyProphet uses semi-supervised learning to calculate a discriminant score from which the p-value of each peak is estimated [8]. It also calculates the false discovery rate (FDR) or q-value (Figure 1) to correct for multiple testing [13].

**Figure 1:**
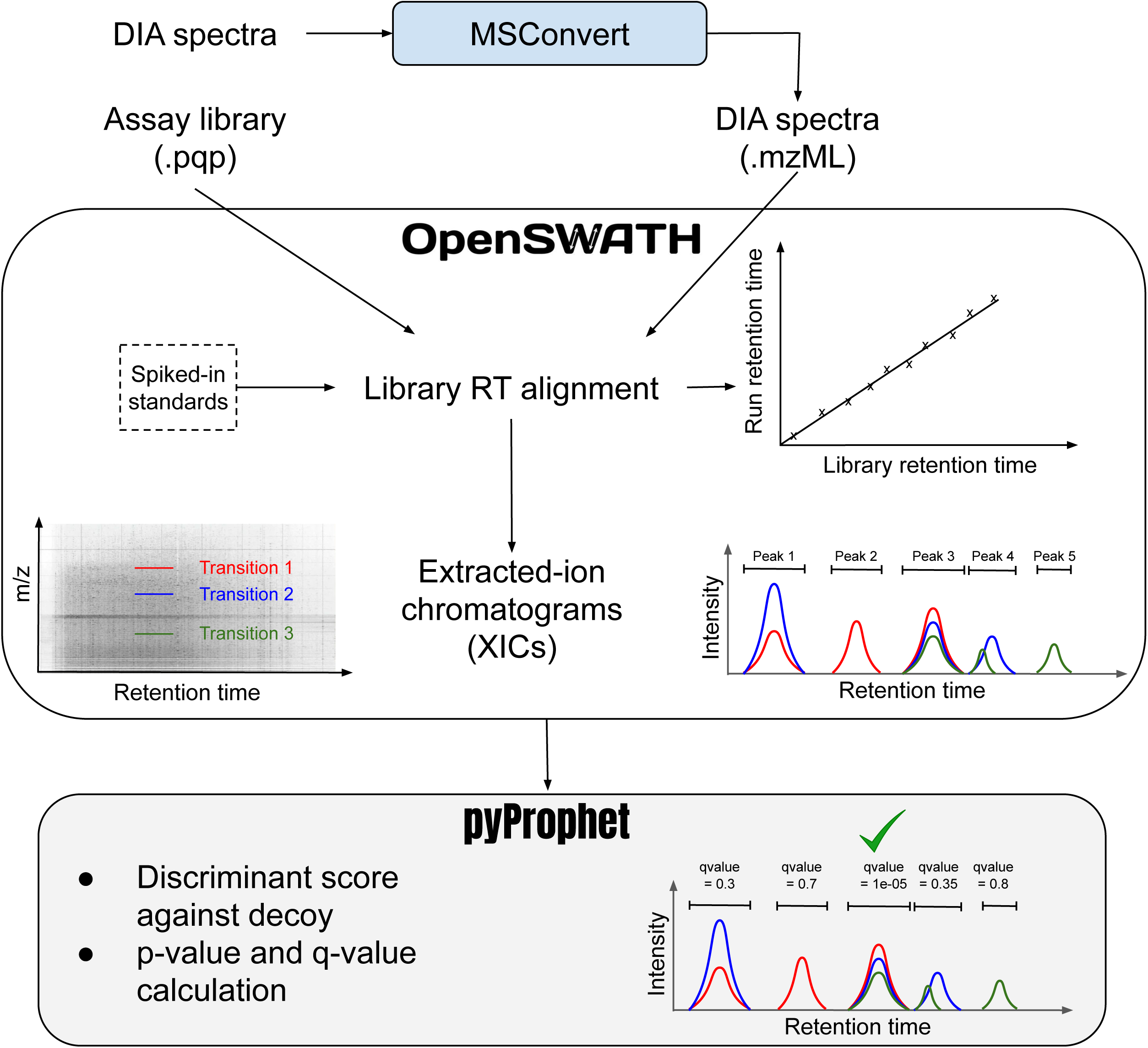
Peak-identification workflow for a peptide precursor with three transitions (red, blue and green) is illustrated. The first step is data conversion to the standard mzML format, subsequently OpenSWATH is used for RT alignment of library to the DIA run. OpenSWATH extracts MS2 chromatograms using library coordinates and identifies potential peaks. Successively, pyProphet identifies correct peaks using statistical scoring and FDR calculation.

OpenSWATH is run independently on each run, which can cause inconsistency in quantification due to altered peak-boundaries, selecting different peaks across runs; and the strict error-rate control by pyProphet may also lead to missed peaks in certain runs [14]. Alignment of MS2 chromatograms across multiple runs establishes consistency and correspondence between the features. For this purpose, a recently developed software DIAlignR is employed: DIAlignR provides highly accurate retention-time alignment across heterogenous SWATH runs [15]. It uses q-values from pyProphet to pick a reference run for each peptide and aligns its XICs to the chromatograms of other runs (Figure 2).

**Figure 2:**
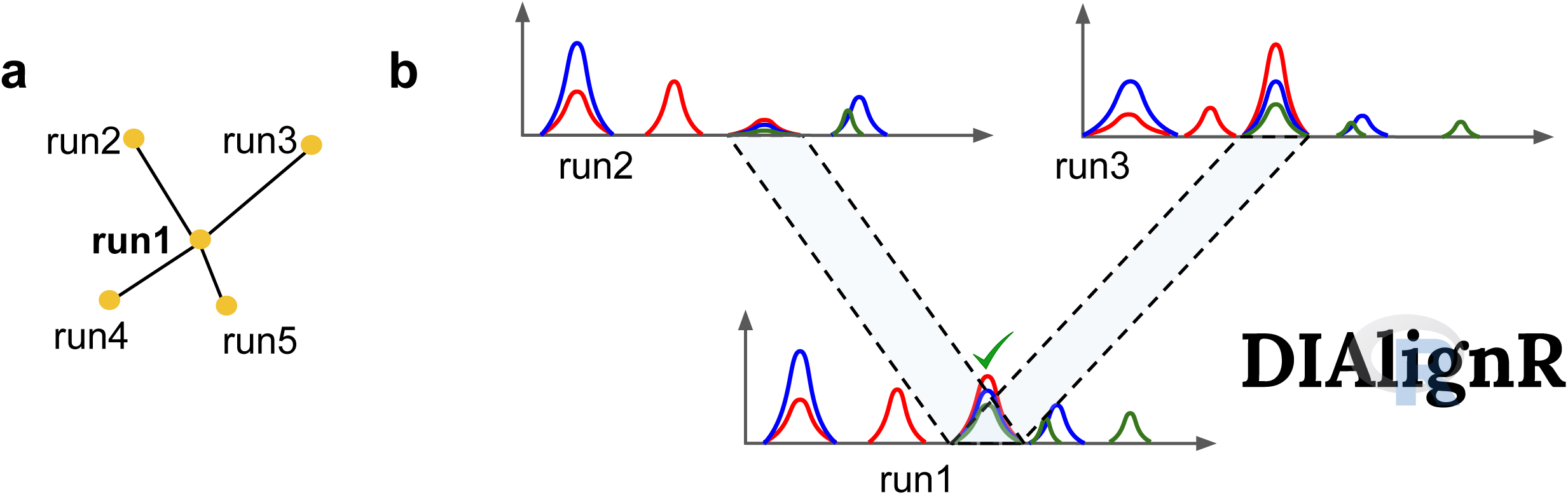
a) For retention-time alignment, DIAlignR picks a reference run for each precursor. b) The XICs of the reference run are aligned to the corresponding XICs of other runs using a non-linear approach. Thus, peak-boundaries from the reference run are mapped to the analysis run, which establishes consistency in quantification. Here, an alignment of XICs of run2 and run3 to a reference run1 is illustrated.

## 2 Materials

To explain the data-analysis workflow an example dataset of *Streptococcus Pyogenes* is used [11, 16]. The source data and intermediate result files can be downloaded from this link: http://www.peptideatlas.org/PASS/PASS01508. Although required applications (OpenMS, pyProphet) are publicly available, we suggest using docker based images for a convenient installation of the applications. For this tutorial, at least 50 GB storage space is required to store spectra and intermediate chromatogram files. The following material is needed to get started:

1. Example dataset: The example dataset PASS01508 has all the intermediary files for this tutorial. Download this repository and name it “PASS01508”. Docker scripts are available at the root level. The “/lib” folder holds assay-library, alignment library, and SWATH acquisition scheme. The “/data” directory has MSConvert output mzML files and raw files inside the “raw” folder. The “/results” folder has OpenSwath output files in the “OpenSwathOutput” folder. It also has an “osw” and an “mzml” directory with features and chromatogram files required for alignment by DIAlignR.
2. SWATH spectra files: A detailed protocol to acquire high-quality SWATH-MS data is provided in Chapter 11. In the example dataset, *S. pyogenes* (strain SF370) was grown in 0% and 10% human plasma in biological duplicates. Four technical replicates for each sample were analyzed on an Eksignet nanoLC (with two-hour linear gradient) coupled to an AB SCIEX TripleTOF 5600 in SWATH-MS mode. Thus, the dataset has a total of 16 runs. The raw files are available at the /data/raw/ directory in PASS01508. The instrument produces two files per run, .wiff and .wiff.scan files, which should always be stored together. Besides the AB SciEx TripleTOF, other instruments can be used to acquire spectra in data independent acquisition mode (*see* **Note 1**).
3. Assay library: In a library-based workflow, the raw file is mined against an assay library which contains precursors sequences, their retention times, their transitions m/z values and intensities. For statistical scoring of peptide identifications, these libraries must contain suitable decoy transition groups [8, 9]. The assay library can be built from one or more shotgun runs [5] or downloaded from the SWATHAtlas database http://www.swathatlas.org. The SpyogenesAssayLibrary_decoy.pqp library used in this workflow is available in the /lib folder at P ASS01508. The conversion to .pqp format is explained at the end of this chapter (*see* **Note 2**). To open the library, either use online SQLite viewer (https://inloop.github.io/sqlite-viewer/) or install a local viewer such as the SQLite DB browser (https://sqlitebrowser.org/dl/).
4. Peptide assays for alignment: For targeted-extraction, the library retention time needs to be aligned to each DIA run using linear or nonlinear methods [17, 18]. Generally, a set of spiked-in standard iRT peptides are used for this, however, in the absence of spike-in standards, the library can be aligned using high-intensity endogenous peptides without sacrificing quantitative accuracy [18]. In the example dataset we have picked 20 endogenous peptides for linear alignment. This assay library is available in the /lib directory at PASS01508.
5. Docker software: Docker isolates dependencies between external software libraries and operating systems (Linux, Mac, Windows), providing reproducible computational results. Since docker images are self-contained, they are lightweight and can be easily ported. First, install the docker engine on your machine. For Linux systems, follow guidelines at https://docs.docker.com/install, for Windows 10 Home install Docker Toolbox from https://docs.docker.com/toolbox/toolbox_install_windows, for Windows 10 Pro or Mac install Docker Desktop from https://docs.docker.com/docker-for-windows. On a Linux setup, use the following commands to get the OpenMS and pyProphet image:

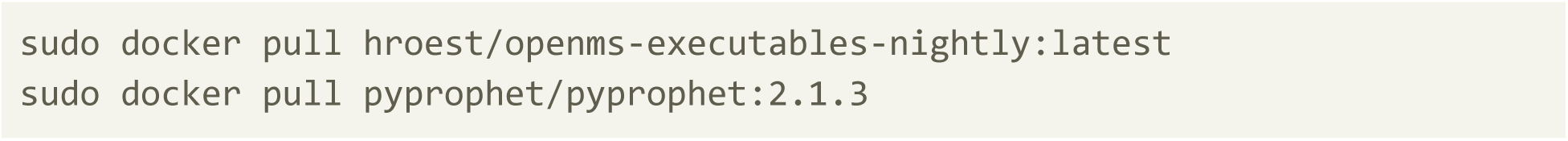 For Windows 10 Home: Click on the start menu and type “Docker Quickstart Terminal”. In this terminal window enter previous commands without *sudo* (*see* **Note 3**)-

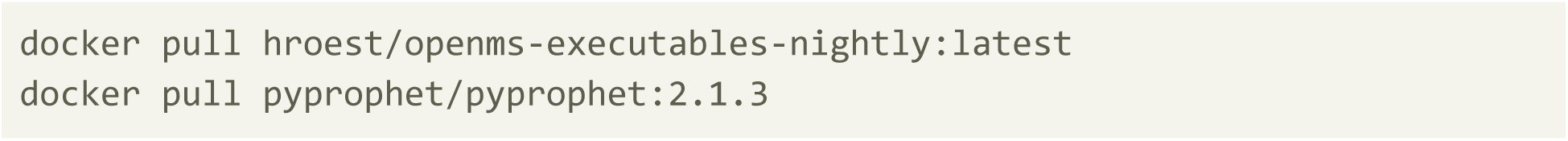
6. Proteowizard: To convert raw files from vendor-specific format to open standardized mzML format, the msconvert tool from Proteowizard software suite is used. This software is available at http://proteowizard.sourceforge.net/download.html. For the software version and getting msconvert for a Linux system see **Note 4**.
7. R and Rstudio: Alignment tool in this tutorial requires R (version > 3.5.0), therefore, install the latest version of R from https://cran.rstudio.com/. Rstudio integrates many tools to visualize data, to document scripts and to work efficiently with R. It can be downloaded from https://rstudio.com/products/rstudio/download/.
8. DIAlignR: This is an R-package which uses raw MS2 chromatograms for alignment of features across multiple runs. It can be installed from Bioconductor in R (R version > 3.5.0). To get this package, open RStudio and in the “Console” type these commands:

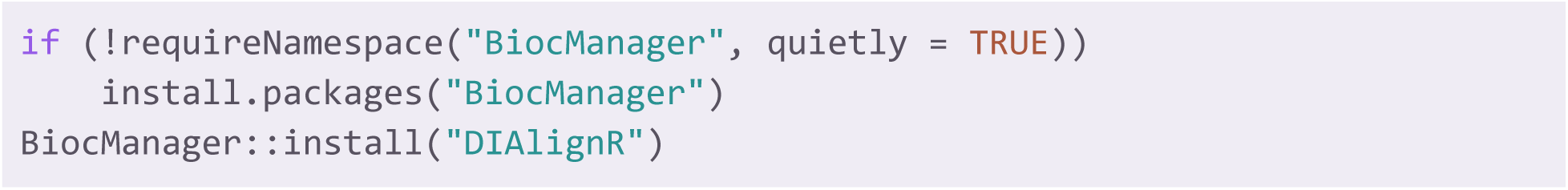

## 3 Methods

Before proceeding to an automated analysis, it is crucial to understand DIA data, quality of chromatography and fragment-ion spectra. The Skyline software provides an open-source interactive solution for data visualization. We recommend using the Skyline-based workflow described previously by Röst et al (2017) to assess data quality and manually inspect few precursors and their transitions [14].

### Step 1: File conversion to mzML

MSConvert, provided with ProteoWizard software suite, is used to convert vendor-specific raw spectra files into an open format such as mzML or mzXML. We will use mzML as it is a standardized format.

1. On a Windows setup, click on the *Start Menu* at the lower left corner (Windows icon). Type “MSConvert” in the *search* field and click on *MSConvert*.
2. Select *List of Files* then click on the *Browse* button and navigate to the folder where the .wiff and .wiff.scan files are present. Select the files and click on the *Add* button. Choose an *Output Directory* using the second *Browse* button. In the *Options* field select following parameters- Output format: mzML; Binary encoding precision: 64-bit; Write Index: yes; Use zlib compression: yes; TPP compatibility: yes; Package in gzip: yes; Use numpress linear compression: yes. Other boxes can be left unchecked (Figure 3). (Optional) To perform centroiding, in the *Filters* menu select *Peak Picking* from drop down; Algorithm: Vendor; MS levels: 1 – 2, click on *Add* and then click *Start*. OpenSWATH produces best results on profile data and should be used for any real-world analysis, however, to reduce file size and subsequent execution time, we chose to use centroiding in this workflow (*see* **Note 5**).

**Figure 3:**
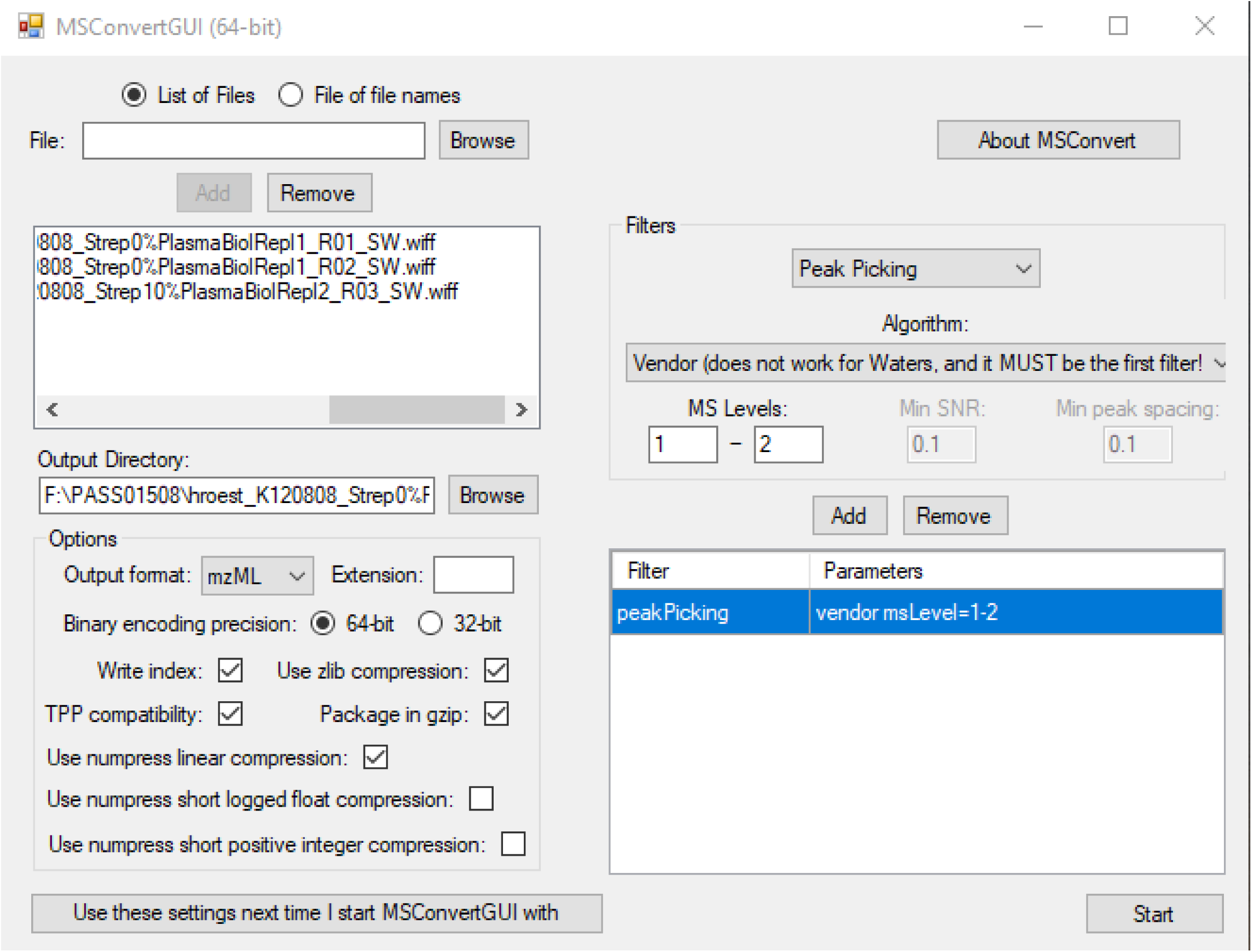
Converting DIA spectra data to mzML format and centroiding spectra using MSConvert GUI

### Step 2: OpenSWATH workflow

*OpenSwathWorkflow* is the main command which efficiently integrates multiple tools from OpenSwath software. A basic command looks like this:

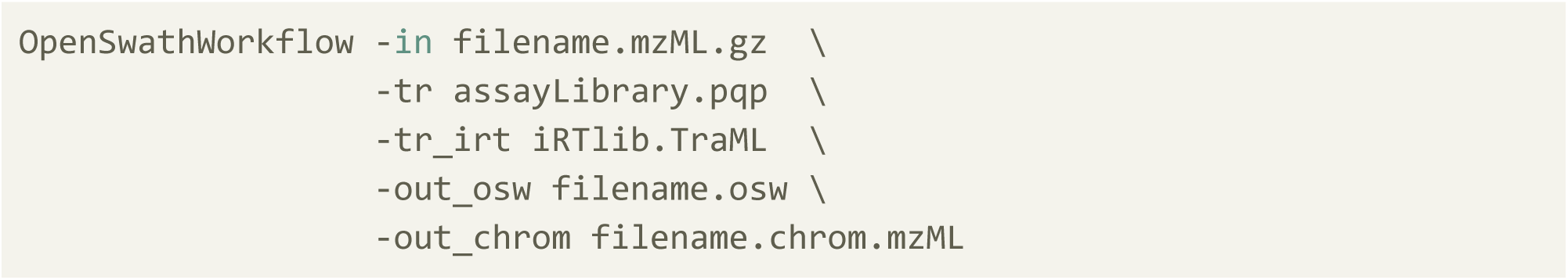

Where -in requires a spectra file, -tr requires an assay library, -tr_irt expects an iRT peptides library, -out_osw expects an output filename that shall contain picked peaks and -out_chrom requires the filename that will contain MS2 chromatograms.

Firstly, move the MSConvert output files to the “data” directory which is located at the root of PASS01508. Make sure the assay library and the iRT library are present in the “lib” directory. For automated analysis-

On a **Linux system**, use the “oswDockerScript.sh” file to run this workflow:

1. Open “Terminal”. In the “Terminal” navigate to the root of PASS01508, where the “oswDockerScript.sh” script is present.
2. Enter the following command at the Terminal:

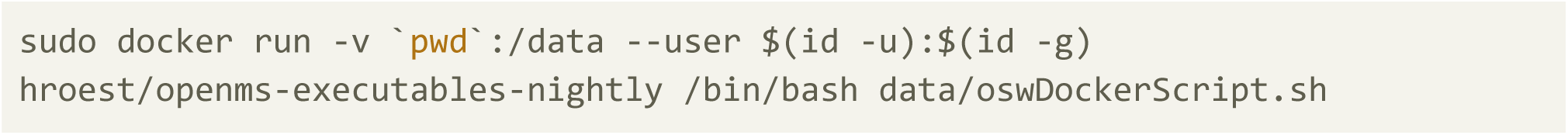

On a **Windows system**, use the “oswDockerScriptWin.sh” file to run this workflow:

1. Click on the start menu and type “Docker Quickstart Terminal”. In the terminal navigate to the root of the data downloaded from PASS01508, where the “oswDockerScriptWin.sh” script is present.
2. Enter this command (*see* **Note 3**):

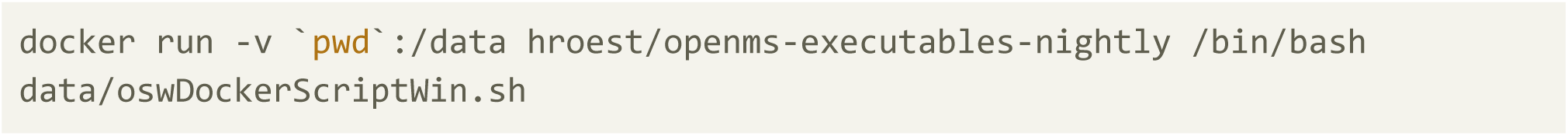

This step yields two files (.osw and .chrom.mzML) for each run. The osw file is a collection of scored peak-groups in SQLite format. This contains potential peaks for each peptide and their scores. The chrom.mzML file contains all XICs of all transitions. For comparison, our results are stored at “results/OpenSwathOutput”.

Besides the flags mentioned above, a few other important flags to control the execution of *OpenSwathWorkflow* are explained in Table 1. These could be added into the docker script used above. The explanation of flags can easily be obtained by following command:

**Table 1:** Few important flags in *OpenSwathWorkflow* command

| Parameter | Explanation |
| --- | --- |
| --help | displays all flags and their explanations. |
| **Isolation window specific** |  |
| -swath_windows_file | A text file specifying the SWATH window out of which the fragment-ion traces will be extracted (see example file). |
| -sort_swath_maps | Sorts table specified in *swath_window_file*. |
| -min_upper_edge_dist | The overlap (in m/z) between neighboring SWATH windows used for data acquisition (Must not be used with *swath_window_file*). Usually set to 1.0 for regular SWATH-MS. |
| **Speed and memory optimization** |  |
| -batchSize | The number of chromatograms that are loaded into memory and analyzed at once. The smaller this number, the less memory required. |
| -readOptions | Sets the memory strategy: by default, OpenSWATH loads all data into memory while “cache” will create a set of cached files on which OpenSWATH will work and not load any data into memory (more memory efficient). A good compromise between these two options is “cacheWorkingInMemory” which uses the cached files but loads the current Swath map into memory. |
| -tempDirectory | Temporary directory to store cached files. (Must be used if *readOptions* are using a caching strategy) |
| -threads | Defines number of threads to parallelize extraction of XICs and their scoring. |
| **Extracted-ion-chromatogram (XIC)** |  |
| -RTNormalization:alignmentMethod | Selects method to align Library RT to sample RT space: ‘linear’, ‘interpolated’, ‘lowess’ and ‘b_spline’. Recommended value with iRT peptides is ‘linear’ and with endogenous peptides select |
|  | 'lowess'. |
| -rt_extraction_window | Size of the RT window in seconds to extract XICs from that peaks should be picked and scored. The RT window is centered around the aligned library RT. Default value is 600 seconds, which means extracting +/- 300 seconds around the expected RT. This value should be changed depending on the quality of library RT alignment. |
| -extra_rt_extraction_window | Adds additional time to the rt_extraction_window only for XIC extraction. Peaks will not be picked and scored from this extra XIC. It is useful for visual inspection of chromatograms and for alignment tools such as DIALignR. |
| -mz_extraction_window | Size of m/z extraction window (in Thomson) centered around library m/z. This value needs to be adjusted according to the instrument resolution. It's unit can be changed with *mz_extraction_window_unit*. |
| -mz_extraction_window_unit | Defines the unit of *mz_extraction_window* and can be set to ppm. |
| **Peak smoothing** |  |
| -Scoring:TransitionGroupPicker:PeakPickerMRM:peak_width | Peak-width in seconds required to define a non-spurious peak (adjust to your chromatography). |
| -Scoring:TransitionGroupPicker:PeakPickerMRM:sgolay_frame_length | Number of points required for convolution by Savitzky-Golay smoothing. Increasing this parameter leads to stronger smoothing. |
| -Scoring:TransitionGroupPicker:PeakPickerMRM:sgolay_polynomial_order | Degree of polynomial fitted by Savitzky-Golay smoothing. Decreasing this parameter leads to stronger smoothing. |
| -Scoring:TransitionGroupPicker:PeakPickerMRM:use_gauss | If set as 'true', Gaussian filter is used for smoothing. |
| -Scoring:TransitionGroupPicker:PeakPickerMRM:gauss_width | Gaussian width in seconds, estimated peak size in seconds. |
| **Debug mode** |  |
| -debug | Set as 1 to print all debug output. |
| **Quantification and Scoring** |  |
| -TransitionGroupPicker:background_subtraction | Set as 'exact' to remove background from peak signal. This option may increase the accuracy of the quantification. |
| -use_ms1_traces | Extract the precursor ion trace(s) and use them for scoring |

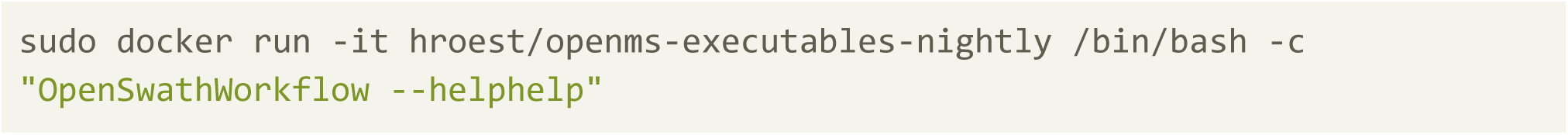

A web-portal http://www.openswath.org/en/latest/docs/openswath.html also goes into additional detail about OpenSWATH. Users are suggested to sign up for the mailing list General OpenMS discussion for OpenSWATH related inquiries.

### Step 3: False Discovery Rate (FDR) by pyProphet

After running OpenSWATH, a q-value for each feature across all runs is calculated using pyProphet. It requires the osw files and the corresponding assay library.

The following steps are performed by pyProphet:

1. Merges all osw files.
2. Trains a classifier on merged.osw. The classifier can be either “LDA” (Linear Discriminant Analysis) or “XGBoost”. To see all flag use the help command:

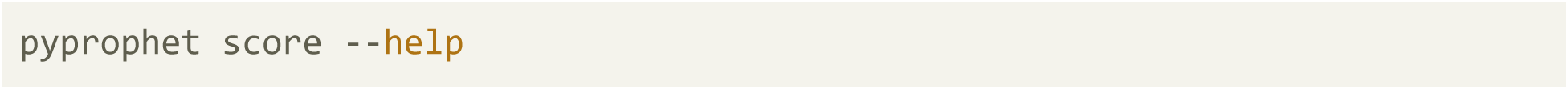
3. Employs (type I) error rate control at peptide level. The FDR-context can be “run-specific”, “experiment-wide” and “global”. To get an explanation of all options use help:

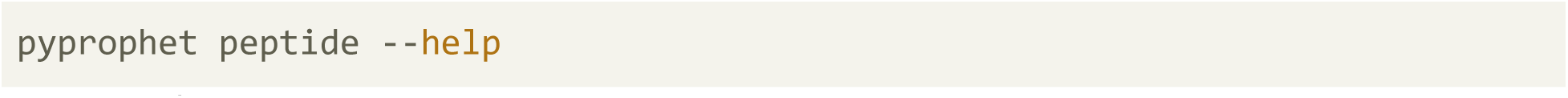

Basic commands for estimating q-value are given below:

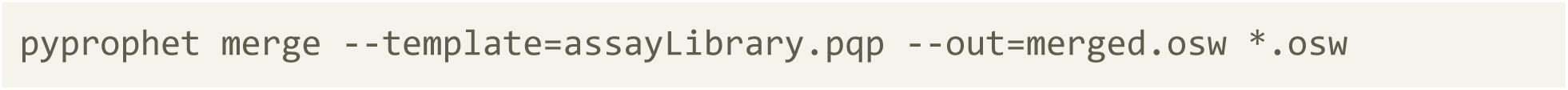

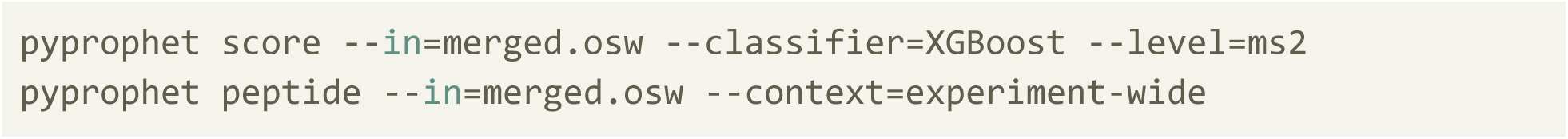

For ease of use, we have included all the commands in a docker script. Before proceeding, make sure osw files from the previous steps are present in the “results” folder.

On a **Linux system**, open the “Terminal”. In the “Terminal” navigate to the root of PASS01508, where the “pyprophetDockerScript.sh” script is present, then, execute following command-

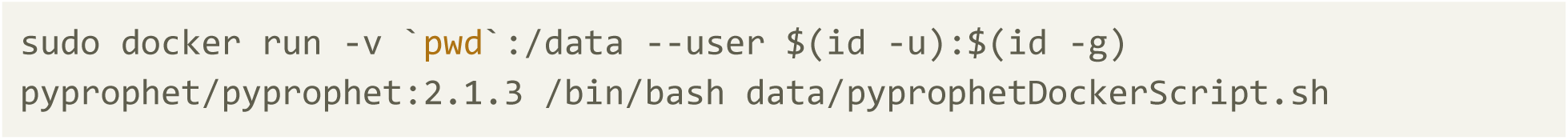

On a **Windows system**, click on the start menu and type “Docker Quickstart Terminal”. In the terminal navigate to the root of PASS01508, where the “pyprophetDockerScriptWin.sh” script is present. Then, enter the following command-

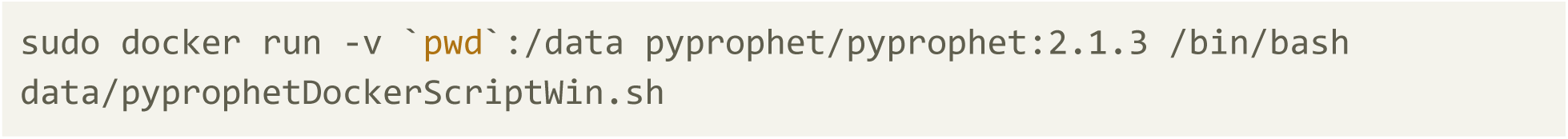

pyProphet outputs a “merged.osw” file that has an FDR value for each analyte; in addition, it also outputs pdf files which contains summary figures on calculated discriminant score (d-score), p-values and q-values. These files must always be consulted so that common errors, such as having an assay library without decoys, library with improper decoys and library from another organism can be identified. In a successful analysis, pyProphet will have a clear separation between positives (true targets) and negatives (false targets and decoys), as shown in Figure 4. The decoy distribution is used to estimate the q-value. To directly get a quantitative matrix from this output, see **Note 6**.

**Figure 4:**
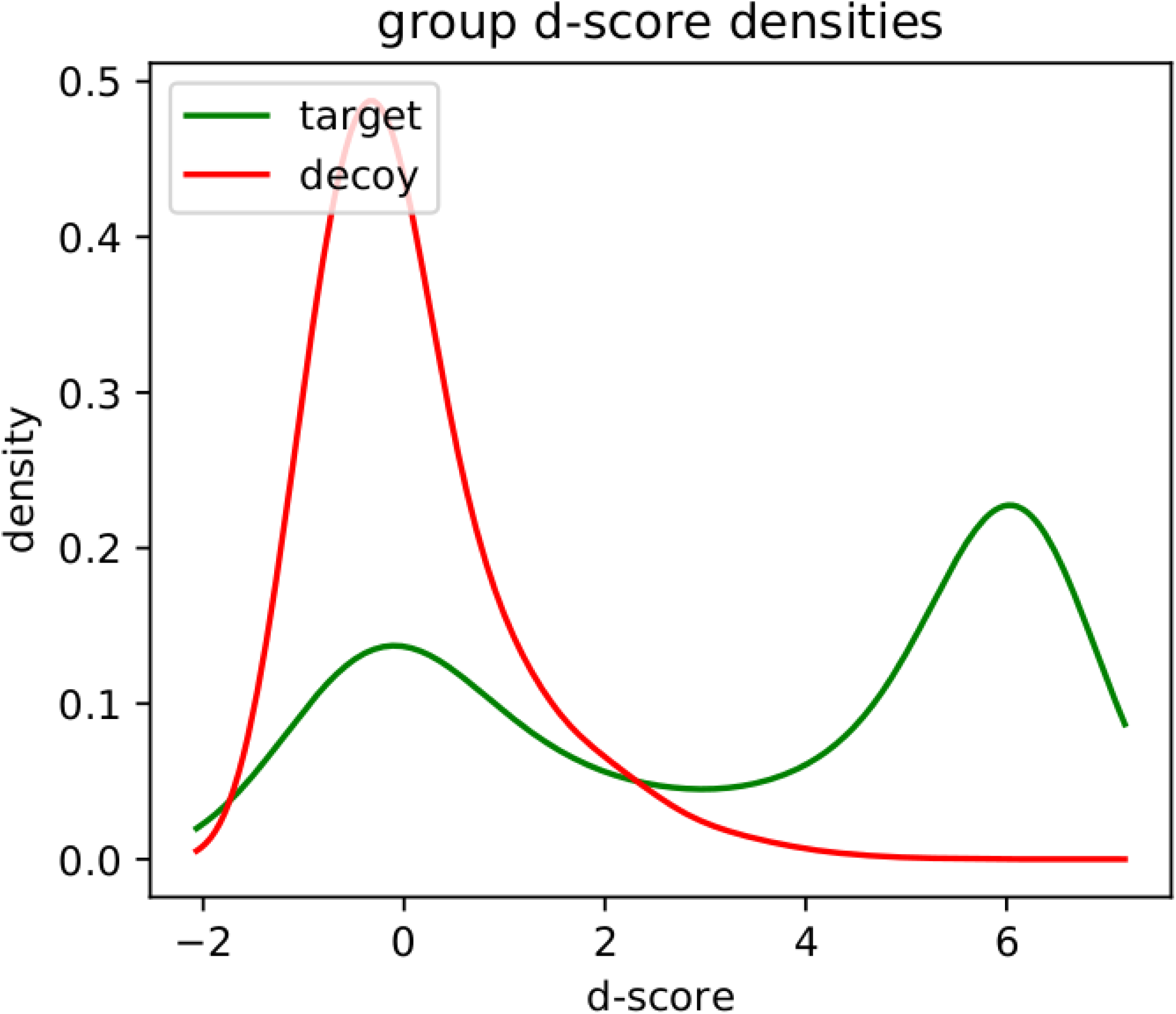
Density-plot from calculated discriminant score (d-score). Generally, a bimodal distribution is expected for target peptides, whereas decoy peptides are expected to have a unimodal distribution. Based on these distributions, pyProphet computes q-values which are used to select a set of results with a well-defined FDR.

### Step 4: Multi-run alignments using DIAlignR

DIAlignR has a set of functions for fetching XICs, selecting reference run for each analyte, performing chromatogram alignment and picking features based on the alignment. A single function *alignTargetedRuns* integrates all the functions and gives an intensity table. Prepare the data by moving the .chrom.mzml files to the “results/mzml” folder and the merged.osw file to the “results/osw” folder. To perform alignment:

1. Open Rstudio.
2. In “Console”, use *setwd* command to navigate to the “PASS01508/results”directory.

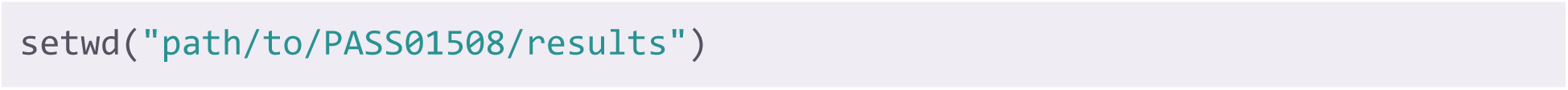
3. To perform alignment and compute the intensity table, use the following command.

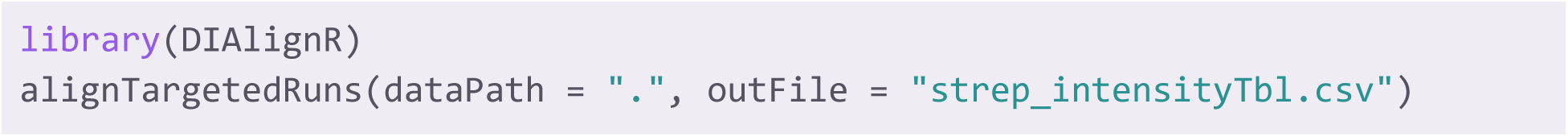

*alignTargetedRuns* has many options to control the alignment algorithm and peak picking. A list of these options can be obtained using `?alignTargetedRuns` in RStudio. Several of these options are explained in Table 2.

**Table 2:**
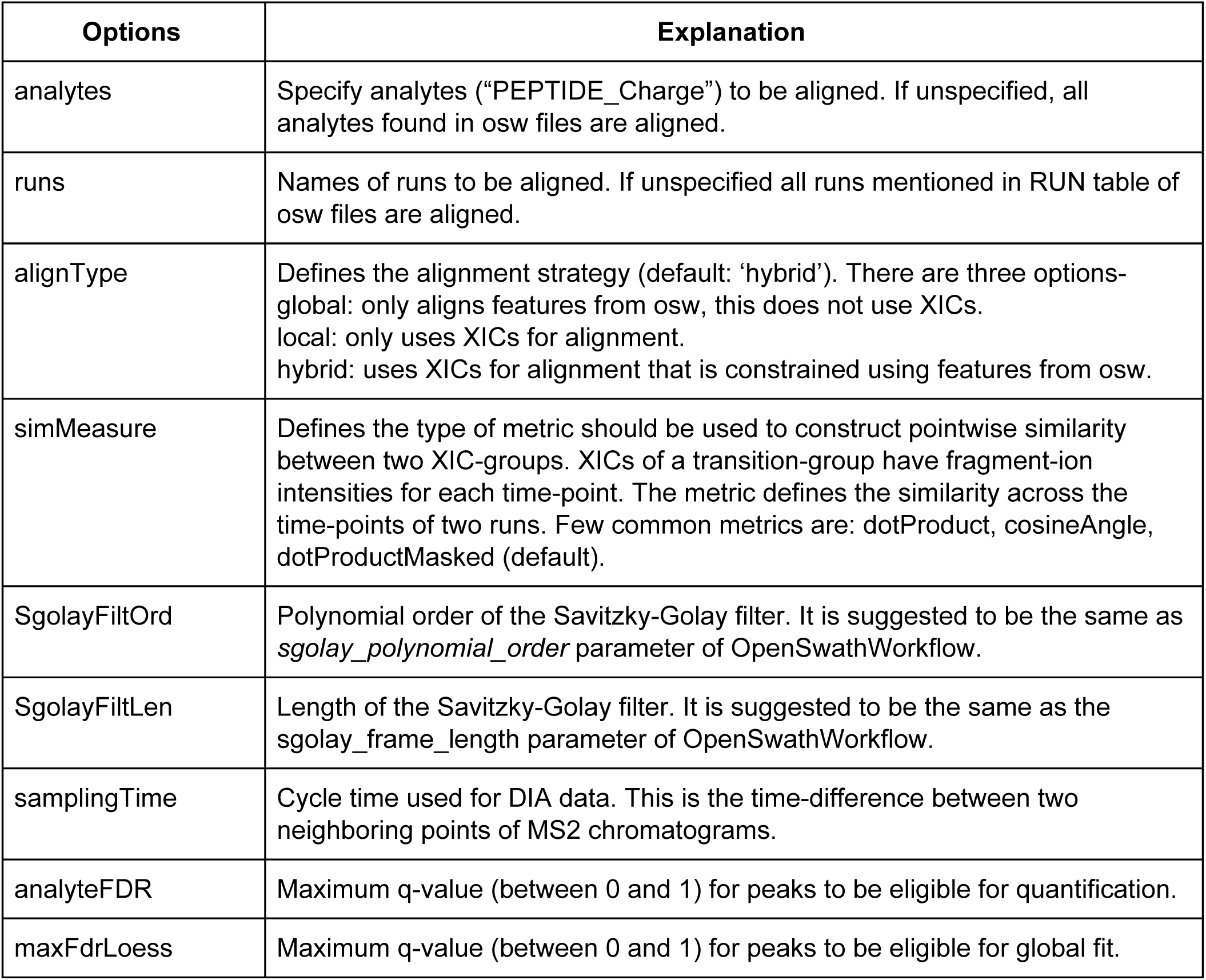
Few options in *alignTargetedRuns* to control alignment of chromatograms

DIAlignR also allows the user to plot aligned chromatograms. Use *getAlignObjs* and *plotAlignedAnalytes* functions to see aligned chromatograms.

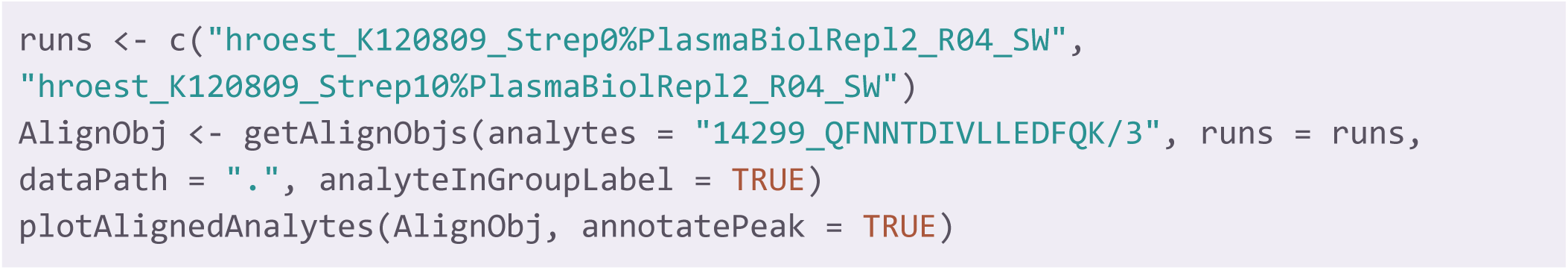

Any issues related to DIAlignR can be reported at https://github.com/shubham1637/DIAlignR/issues.

## 4 Notes

1. SWATH-MS was originally developed for a TOF based mass-analyzer using SCIEX instrumentation. However, SWATH-like data-independent acquisition (DIA) data can be obtained from other types of instruments. One of the major considerations is the acquisition speed, which needs to allow sufficient sampling during elution of an analyte from the LC column. OpenSWATH can analyze data from multiple vendors, including Waters, Thermo Fisher and Bruker. OpenSWATH, Skyline and other software have been used to analyze DIA data from Thermo Q Exactive instruments [19-22].
2. While OpenSWATH supports the .TraML format, to reduce the downstream file size, the SQLite based .pqp format is recommended. To convert a library from TraML to pqp format, open “Terminal” (in Linux) or “Docker Quickstart Terminal” (in Windows), as explained in the Methods section. Navigate to the location where the .TraML library is present, then run the following command:

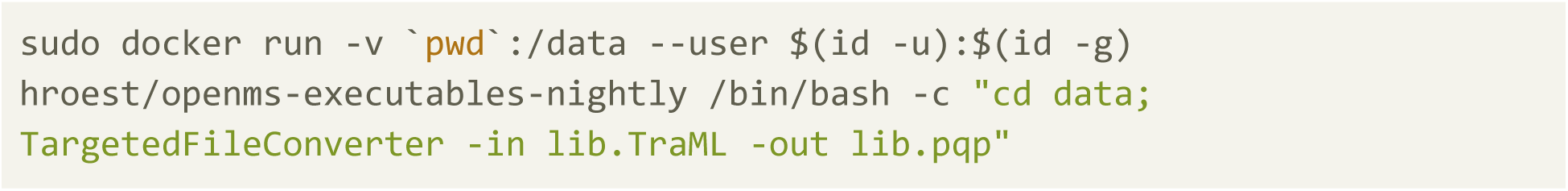
3. To execute docker commands for OpenSwath and pyProphet on a Windows 10 Home setup, the following precautions must be taken:
  - To run docker containers in Windows and Mac systems, Linux commands can be used without having leading *sudo*.
  - Make sure there are no spaces in the file path.
  - Make sure PASS01508 data is in the C: drive (shared drive are not enabled for docker by default).
4. In this tutorial MSConvert is used from ProteoWizard release: 3.0.11252 (2017-8-22). For Linux systems, the docker image can be downloaded using the following command:

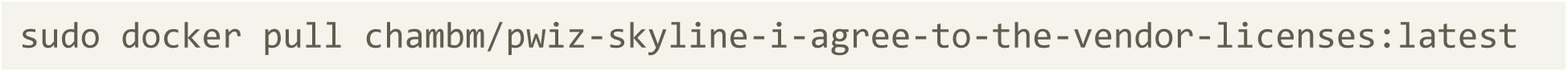 To convert wiff files to mzML, use commands from “msconvertOnLinux.sh” file available at P ASS01508.
5. OpenSWATH produces optimal results on profile data. To obtain profile data, do not specify anything in the *Filters* menu and remove any filter from the lower-right box, if present (Figure 3). Centroiding reduces the number of peaks in the data and sometimes may remove low-intensity peaks, the XICs generated by OpenSWATH become less smooth and peak detection becomes less sensitive. If centroiding is performed, it is crucial to compare the results of OpenSWATH to those obtained on profile data on the same dataset to obtain an accurate estimation of how centroiding affects the identification rate [14].
6. Alignment is a necessary step for consistent analysis across multiple runs. Nonetheless, this is a time-consuming step and under some circumstances is not required. A (un-aligned) peptide data-matrix can be directly obtained from merged.osw file, using the following command:

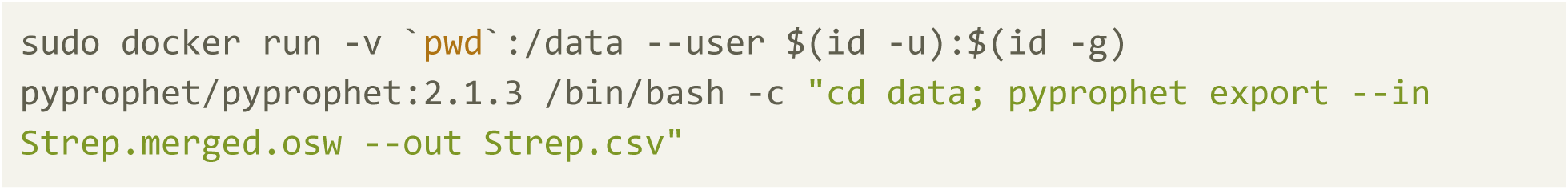

## Notes

http://www.peptideatlas.org/PASS/PASS01508

### Competing Interest Statement

The authors have declared no competing interest.

### Summary of Updates

Code for DIAlignR is updated

